# New Curcuphenol Analogues Possess Anti-Metastatic Biological Activity

**DOI:** 10.1101/2023.05.15.540833

**Authors:** Samantha L.S. Ellis, Lilian L. Nohara, Sarah Dada, Iryna Saranchova, Lonna Munro, Kyung Bok Choi, Emmanuel Garrovillas, Cheryl G. Pfeifer, David E. Williams, Ping Cheng, Raymond J. Andersen, Wilfred A. Jefferies

**Author notes:** Denotes co-first authorship (Samantha L.S. Ellis, Lilian L. Nohara). Corresponding and Senior Author.

## Abstract

For eons, turmeric and curcumin have been used as culinary spices and as traditional medicines and as vogue dietary supplements for a growing list of disorders, including arthritis, digestive disorders, respiratory infections, allergies, liver disease, depression and cancer. The activities of these spices are commonly attributed to curcuminoids; however, the medical applications of this class of compounds has been limited due to the low water solubility, chemical instability, acid lability, poor absorption, rapid catabolism by enzymes of the diverse curcuminoids contained in turmeric and curcumin extracts. Furthermore, identifying the bio-active curcuminoids with unique molecular entities responsible for specific medicinal benefit is at its infancy. To overcome these many issues and substantially advance this area of inquiry, we created a water-soluble achiral curcuphenol analogue and a water-soluble racemic analogue that have enhanced chemical characteristics and biological performance, and we subsequently demonstrated their ability to reverse the immune-escape phenotype, a process that enables tumours to hide from host immune responses and thereby provides tumours a significant growth advantage to metastatic tumours. The discovery that curcuphenols can reverse tumour immune-escape mechanisms and thereby reduce tumour growth, provides a rationale for the development of advanced dissecting nutraceuticals and bioceuticals for unique chemical entities as therapeutic building blocks to synthesize analogues with optimal chemical characteristics capable of harnessing the power of the immune system to extinguish metastatic cancers and beyond.

## Introduction

It is estimated that over the past century approximately 50% of all new anti-cancer drugs including paclitaxel, vincristine, doxorubicin, and bleomycin (1, 2) were derived from natural products (3). Foods including spices and herbs have long been considered to possess medicinal properties. The progress to reduce the many subjective biological activities of these compounds, often classed as nutraceutical or bioceutical, to specific chemical constituents to their unique bio-active molecular entities possessing specific medicinal benefit, has thus fair been largely unsuccessful. For example, extracts of turmeric and curcumin have been purported as natural products that exhibits antioxidant, anti-inflammatory, anti-microbial, anti-parasitic, and anti-cancer activities. However, the low water solubility, chemical instability, acid lability, rapid catabolism by enzymes in the human body, all limit the bioavailability of curcuminoids (4). Additionally, adding to these problems, only a small fraction of ingested curcuminoids contained in turmeric and curcumin extracts becomes absorbed into the bloodstream (4). Attempts have been made to overcome some of these issues by creating new formulations of curcuminoids such as, by creating edible microparticles or nanoparticles (4). In addition an added complexity is the chemical diversity of curcuminoid molecules in turmeric and curcumin. Moreover, comparison between research studies with differently sources of curcuminoid extracts, is virtually impossible as there exists no batching standardization between studies. Thus, despite 100s if not 1000s of years of recorded usage in traditional medicines, translating the use of curcuminoid to clinical practice in cancer, for example, becomes more achievable if individual curcuminoids are identified, with specific mechanisms of action.

Metastatic cancer accounts for 90% of all cancer-related fatalities., it is clearly of the great importance to understand the mechanisms that facilitate the transition from primary cancers to metastatic sequela (5). The cellular immune system has been demonstrated to play a crucial role in cancer progression through the recognition of cancer cells via the antigen processing pathway (APM) (6, 7, 8, 9, 10, 11). This is achieved via the catabolism of tumour-associated proteins into small peptides and the subsequent binding of peptides to major histocompatibility class I molecules (MHC-I) that present antigens to cytotoxic T lymphocytes (CTLs) (6, 8, 10, 11, 12, 13, 14). For the generation of these peptides, endogenous proteins are proteolyzed via the cytosolic proteasome before being transported to the endoplasmic reticulum (ER) by transporters associated with antigen processing 1 and 2 (TAP-1/2) (7, 10, 15, 16, 17). In the endoplasmic reticulum, peptides are bound to MHC-I molecules prior to being carried to the cell surface. (6, 7, 8, 10, 12, 13, 18). When CTLs interact with cell surface MHC-I peptide complexes, they discriminate between normal, cancer, or pathogen-infected cells (6, 7, 8, 12, 13). This interaction is followed by the initiation of an appropriate immune response, often destroying cancer cells or cells infected with pathogens (7, 19, 20).

During metastatic cancer evolution, several genetic and epigenetic alterations, known as metastatic signatures, can occur that allow cancers to escape immune recognition and therefore to continue to grow unabated. There are many mechanisms underlying escape from immune recognition, acting alone or in combination including tumour-induced T-cell anergy, the absence or low expression of MHC-I molecules (21, 22), or defects in the MHC-I antigen presentation machinery (23, 24, 25) and finally in the newly described IL-33 autocrine mechanism that facilitates normal MHC-I expression levels in normal epithelial cells and primary tumours that is defective in many types of metastatic tumours (13). Furthermore there is an established correlation between MHC down-regulation and poor prognosis, including lung cancer (26, 27, 28), breast cancer (29, 30), renal carcinoma (31), melanoma (32, 33), colorectal carcinoma (34), head and neck squamous cell cancer (35), cervical cancer (36), and finally in prostate carcinoma(37). Finally, other than HDACi (12, 38, 39), and the recently described effect of cannabinoids (40) very few small molecules have been described that revert MHC-loss in metastatic tumours have been described. Molecules possess the ability reverse immune-escape have the potential to become mainstream drugs for metastatic disease.

Here we focus on the anti-cancer effects of curcuphenol, a fascinating curcuminoid found to occur natural in spices such as turmeric and curcumin. These studies were initiated because we recently screened a library of marine natural products and identified curcuphenol as a compound that is able to reverse MHC-loss *in vitro* (41). Here we provide chemical approaches to overcome many of the afore mentioned issues disabling the applications of curcuminoids and undertook the total chemical synthesis of novel water-soluble curcuphenol analogues and explored their role in enhancing immune recognition resulting in the reduction of the growth of metastatic tumours in murine mammalian models.

## Results

### Description of TC-1 and A9 Cell Lines

The metastatic A9 cell line is derived from the mouse lung cancer cell line TC-1, which itself originates from primary lung epithelial cells of a C57BL/6 mouse. The immortalization of TC-1 was achieved using the LXSN16 amphitropic retrovirus vector containing Human Papillomavirus E6/E7 oncogenes and subsequently transformed with a pVEJB plasmid expressing the activated human H-Ras oncogene. The A9 cell line is a result of in vivo immunization procedures in animals carrying the original TC-1 parental cells, promoting the selection of clones with improved immune resistance. Unlike the TC-1 parent cells, which exhibit high TAP 1 and MHC I expression, A9 cells display nearly untraceable levels of MHC I. A9 cells are cultivated in Dulbecco’s Modified Eagle Medium (DMEM; Gibco) with 10% fetal bovine serum (FBS; Gibco), 100 U/mL penicillin-streptomycin (Gibco), and kept at 37 °C in a humidified atmosphere with 5% CO2.The A9 murine metastatic lung cancer cell line was chosen to assess as it is known to have reduced expression of APM and to have undergone immune-escape or immune-editing (12, 38, 39). The metastatic A9 cell line is derived from a primary murine lung carcinoma TC-1 that maintains APM expression during in *vivo* passaging.

### A Curcuminoid that induce the antigen presentation in metastatic tumours

In the past, our lab employed Cellomics and Flow Cytometry to examine marine invertebrate extracts gathered from global oceans for their capacity to stimulate TAP-1 and MHC-I expression in the A9 metastatic cell line (41).The active ingredient was identified by NMR and optical rotation as S- (+)-curcuphenol. MHC Class I expression is higher in TC1 primary cell lines than A9 metastatic cell lines, as was anticipated by our previous research. Curcuphenol was purified from sponge extracts as described in (42) Curcuphenol (**Figure 1A**) was observed to increase the abundance of surface expressing of MHC class I protein in metastatic A9 cells, as demonstrated by FACS. An ideal concentration to promote increased expression of MHC class I, but not kill the A9 cells was found to be 12.00815 ug (**Figure 1A**). Optimum concentration to balance MHC class I upregulation and viability was 0.364 ug (**Figure 1A**). MHC class I protein surface expression was observed to be slightly higher on TC-1 cells than curcuphenol induced A9 cells, as determined by the higher APC fluorescence using flow cytometry (**Figure 1B**).

**Figure 1.**
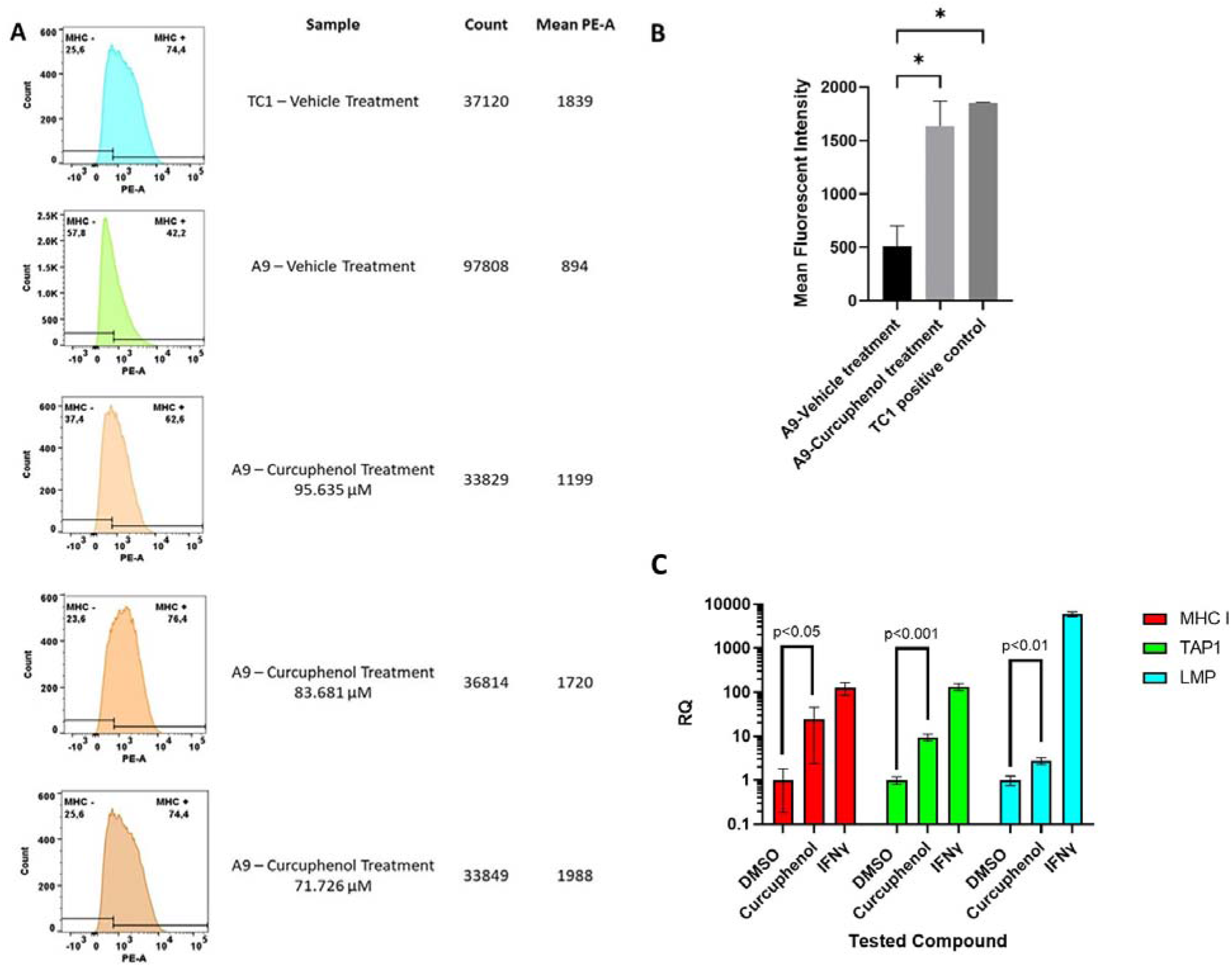
Curcuphenol treatment enhances surface MHC class I expression. **A)** Metastatic A9 tumour cells were treated with either vehicle (DMSO) or curcuphenol for 48 hours. The cells were then harvested and subjected to flow cytometry (The experiment was repeated three times). **B)** The statistical test was done using paired T test. Primary TC-1 tumours cells that constitutively express MHC I were used as a positive control. The mean fluorescent intensity of Metastatic A9 tumour cells untreated and the DMSO treatment was pooled as control population. The values of three Curcuphenol treatments of Metastatic A9 tumour cells were pooled together for the treated group. A two-tailed T test was conducted with a cut-off of p value of 0.05 and below considered significant. The paired T test resulted in a p value of 0.01 which being less than 0.05 is considered significant. This experiment was repeated 3X. **C)** Curcuphenol increased mRNA expression of MHC I, TAP1, and LMP, in A9 cells. RNA was isolated and reverse transcribed into cDNA for Real-Time PCR analysis. Samples were run in triplicate (n = 3) on a 7500 Fast Real-Time PCR system (Applied Biosystems). Relative Quantification (RQ) was calculated by the 7500 Fast Real-Time PCR system software. Values were normalized to Housekeeping gene SDHA106 and calculated with 1% DMSO treatment as reference. Data was visualized and subjected to an unpaired t-test for statistical analysis using GraphPad Prism 9.

IFN gamma (IFNγ) was used as a positive control, inducing a high amount of MHC class I protein expression. There was a significant increase in MHC class I protein expression in A9 cells upon treatment of curcuphenol (**Figure 1B**). Finally, TAP and MHC class I mRNA appeared to increase upon these treatments with optimum concentrations of curcuphenol (**Figure 1C**). Compared to the vehicle control (1% DMSO), stimulation with Curcuphenol appears to increase expression of the antigen-presenting molecule (APM), MHC I, TAP1, and LMP, with an RQ of 24.2 (p<0.05), 9.48 (p<0.001), and 2.77 (p<0.01), respectively (**Figure 1C**).

### Induction by Curcuphenols of T Lymphocyte recognition of Metastatic Tumours

Carboxyfluorescein succinimidyl ester (**CFSE**) is a cell permeable fluorescent dye that covalently coupled to intracellular molecules and can therefore be retained within cells and dilute in fluorescent intensity as the cells divide. CFSE experiments were conducted using CD8+ T lymphocytes from SIINFEKL- specific OT1 mice to determine if curcuphenol can induce antigen presentation of metastatic A9 Cells. Three groups were used to stimulated OT1 T lymphocytes recognition: A9 cells pulsed with SIINFEKL (Vehicle Control); A9 cells induced with IFNγ and pulsed with SIINFEKL (IFNγ T cells); A9 cells induced with curcuphenol and pulsed with SIINFEKL (curcuphenol). The negative control consists of CD8+ T lymphocytes from SIINFEKL-specific OT1 alone (negative CTRL). Evidently, IFNγ and curcuphenol-stimulated CD8+ T cells from SIINFEKL-specific OT1 exhibited more proliferation than the controls. To compare cell proliferation statistically among different treatment groups, the total cell number from various treatment groups at each generation was plotted and assessed using a one-way ANOVA with Tukey’s multiple comparison test. P values less than 0.05 were deemed significant. At generation 1, the IFNγ and curcuphenol-treated cells did not show significantly higher proliferation compared to the vehicle-treated group, whereas at generation 2, the IFNγ and curcuphenol-treated groups displayed significantly greater proliferation than the vehicle-treated group (Figure 2). Overall, these findings support the conclusion that Curcuphenol can reverse the MHC-I immune-escape phenotype in metastatic tumors, leading to immune recognition of tumors by CD8+ T lymphocytes.

**Figure 2:**
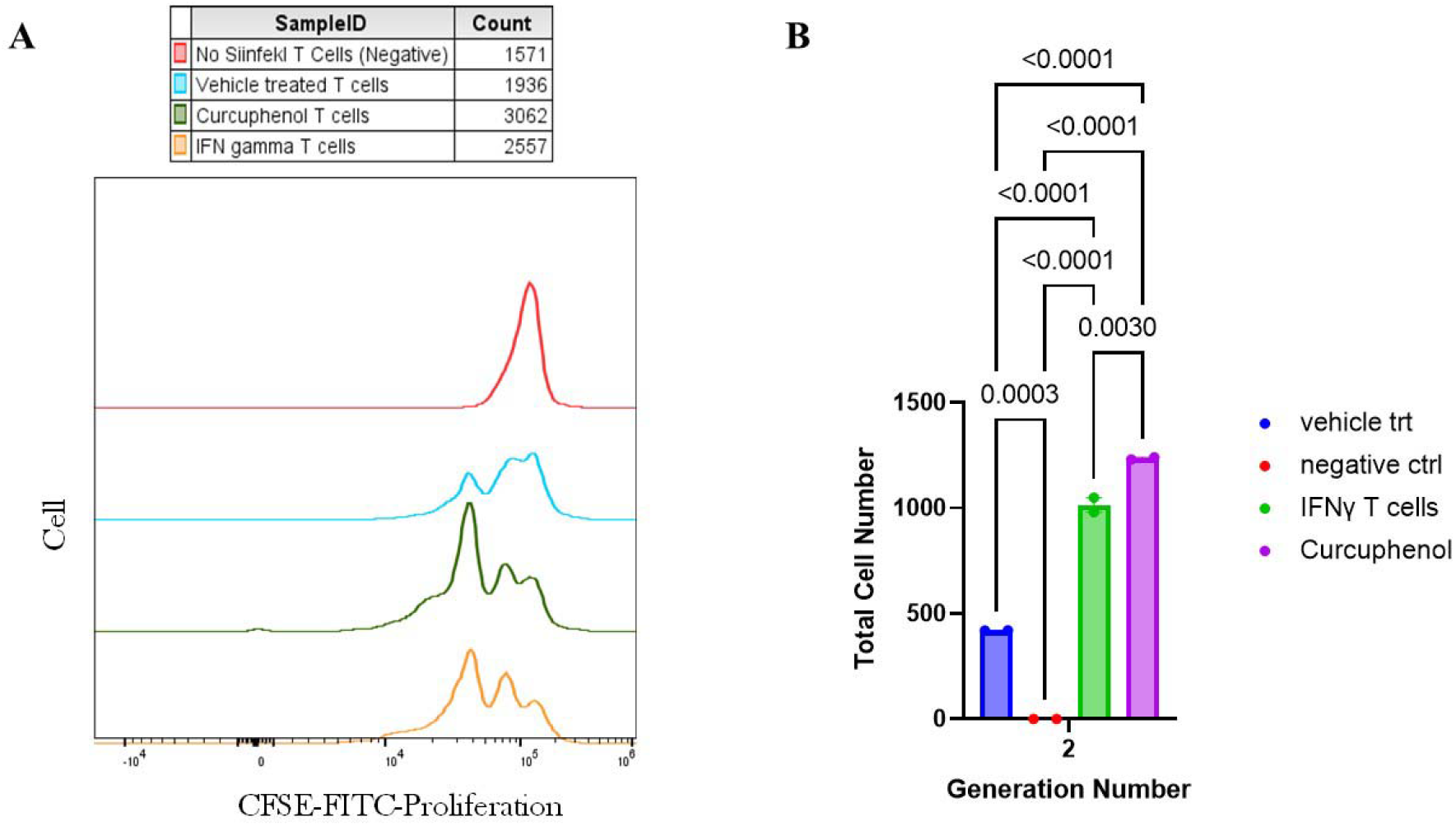
Measurement of Antigen Presentation by Curcuphenol Treated Metastatic Lung tumours. The evaluation of antigen presentation by Curcuphenol-treated metastatic A9 tumors in an OT-I CFSE proliferation assay involved using CD8+ T cells from SIINFEKL-specific OT1 mice to respond to the SIINFEKL peptide presented on MHC class I of A9 cells. A9 cells were treated with 0.00167 µmol of Curcuphenol or 5.832×10-6 nmol/mL of IFN gamma as a positive control. The negative control consisted of OT1 cells alone labeled with CFSE proliferation dye, which decreases within the daughter cells as the generations progress. At generation 1, there was no significant difference in proliferation between the Curcuphenol PO3 analog-treated cells and the uninduced vehicle (peptide) treated group. However, from generation 2 onward, the Curcuphenol-treated group exhibited significantly higher proliferation compared to the uninduced vehicle (peptide) treated group or the CFSE-labeled OT1 T cells alone.

### Synthesis of racemic curcuphenol and structural analogues

Curcuphenol was isolated in the pure S enantiomer from sponge extracts. The synthesis of pure S curcuphenol in the laboratory is challenging. Therefore, we chose to undertake synthesis of a small library of curcuphenols analogues that either lack the chiral centre, PC-02-113 (**11**) or are racemic mixtures to see if we could find an analogue that had better activity *in vivo*. The synthetic methods used to make the analogues as well as racemic curcuphenol as a reference standard is described in the methodology section and highlighted in Figure 3A. The structures and the SwissADME Output for TSA and Curcuphenol and P02-113 and P03-97-1 Analogues is shown in Table 1.

**Figure 3.**
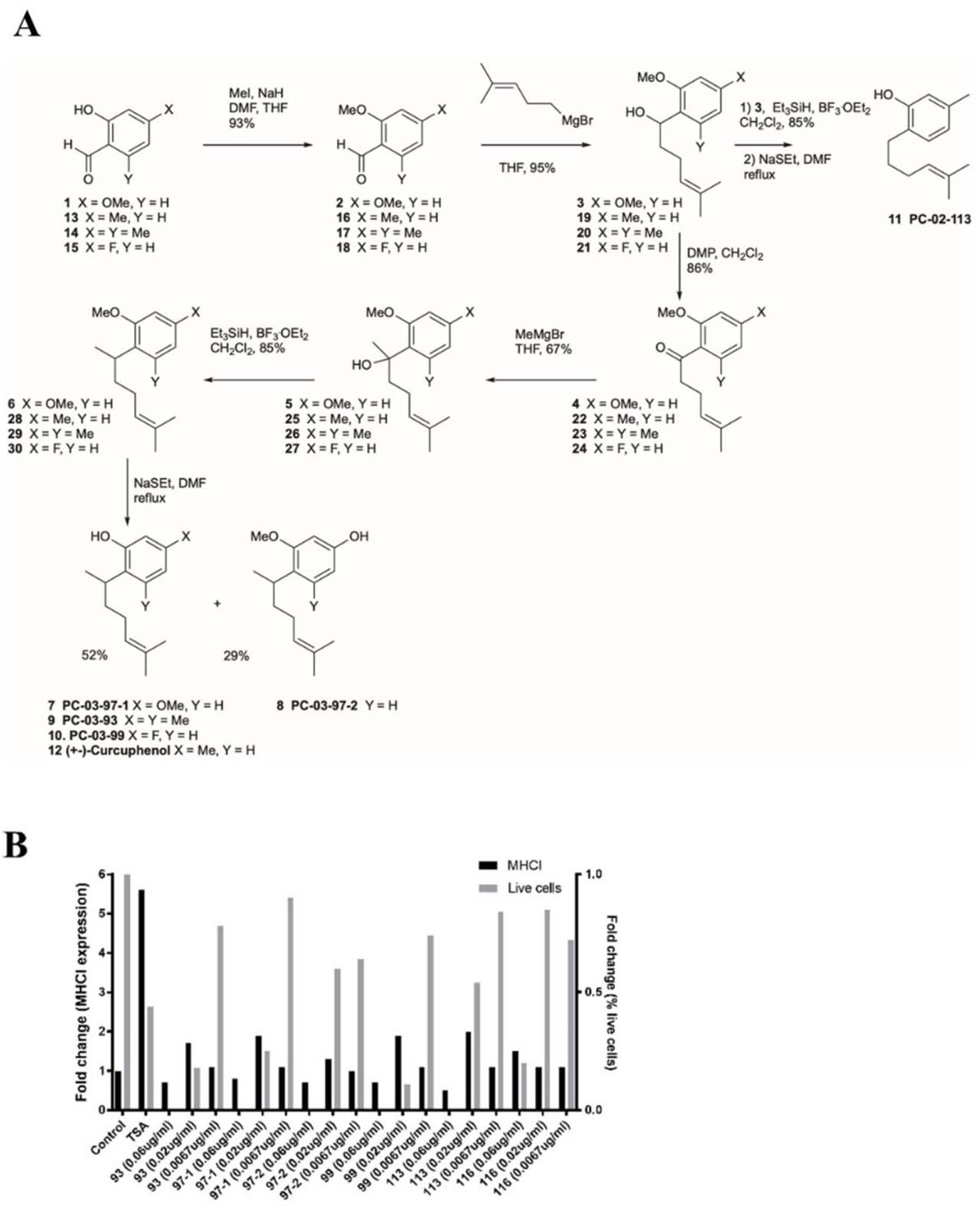
**A.** The process for the synthesis of curcuphenol analogs. **B.** The capacity of P02 and P03 curcuphenol analogs to stimulate MHC-I expression was evaluated using flow cytometry. The black bars refer to the expression of the protein complex, MHC I, on the cell surface, as assessed by flow cytometry. The grey bars refer to the presence of live cells in the sample as a read-out of drug toxicity, as assessed using the Cellomics machine. The experiments were repeated 2 times (n=2) to accurately calculate the significance of MHC-I expression.

**TABLE 1.**
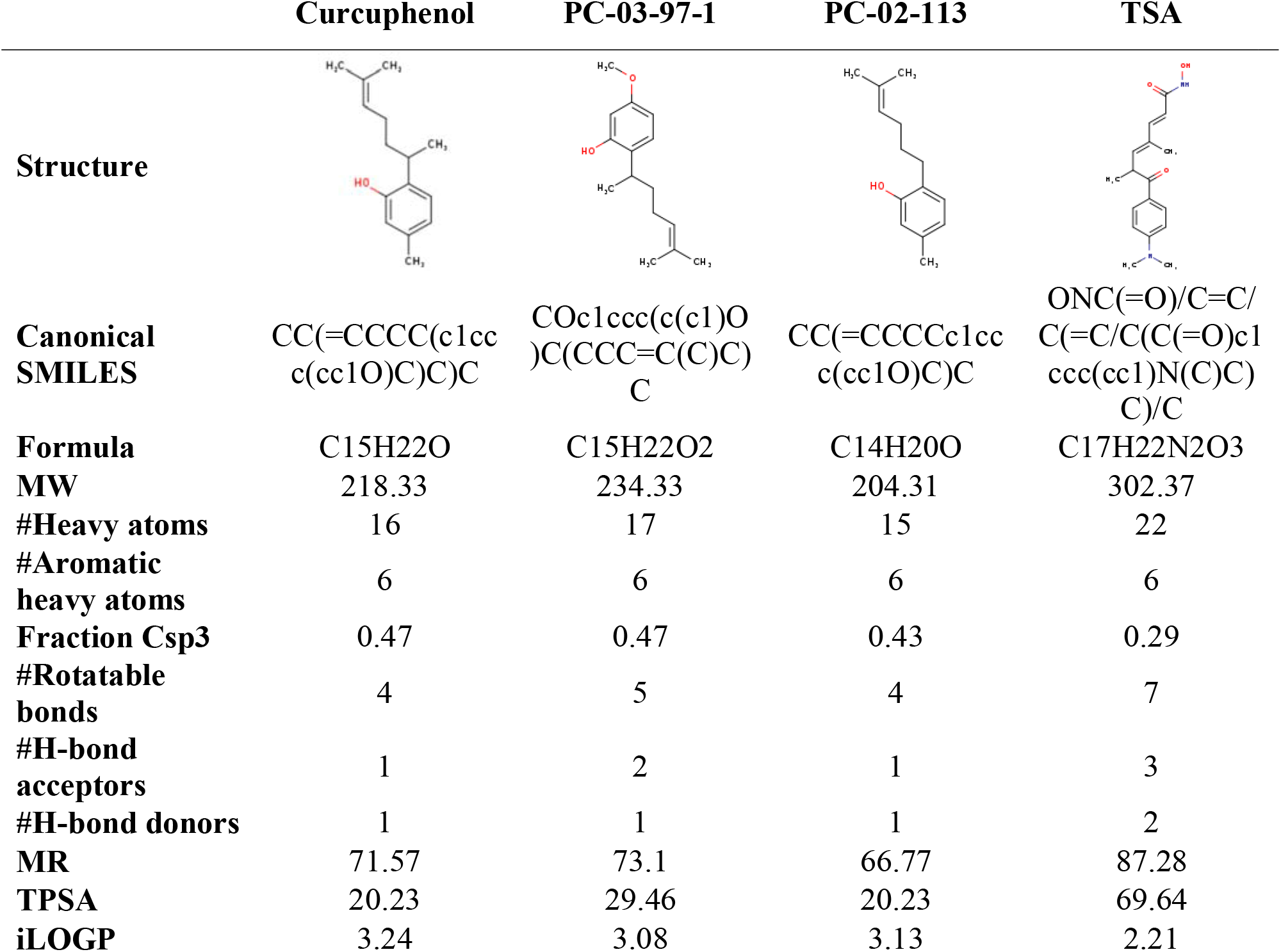
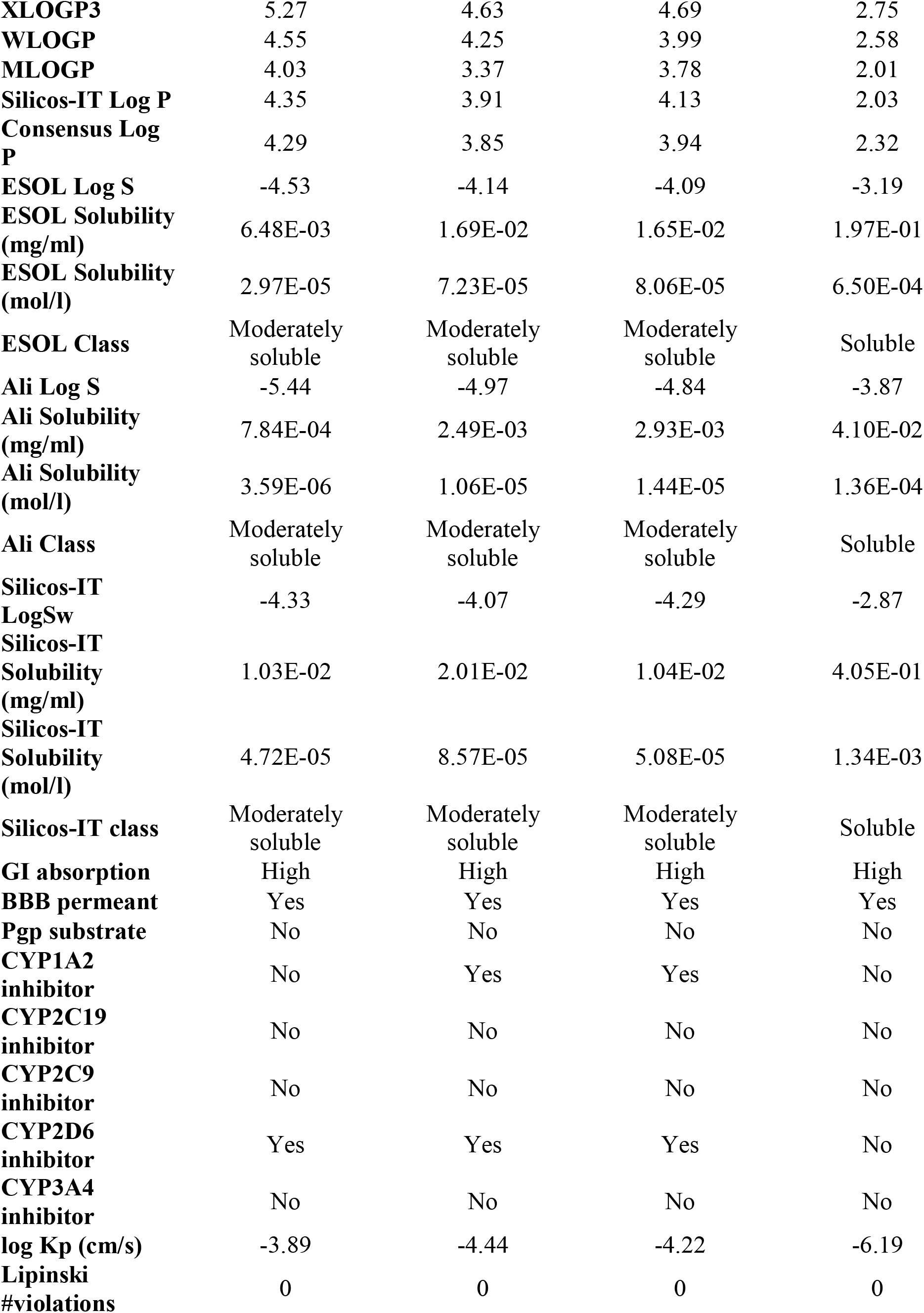
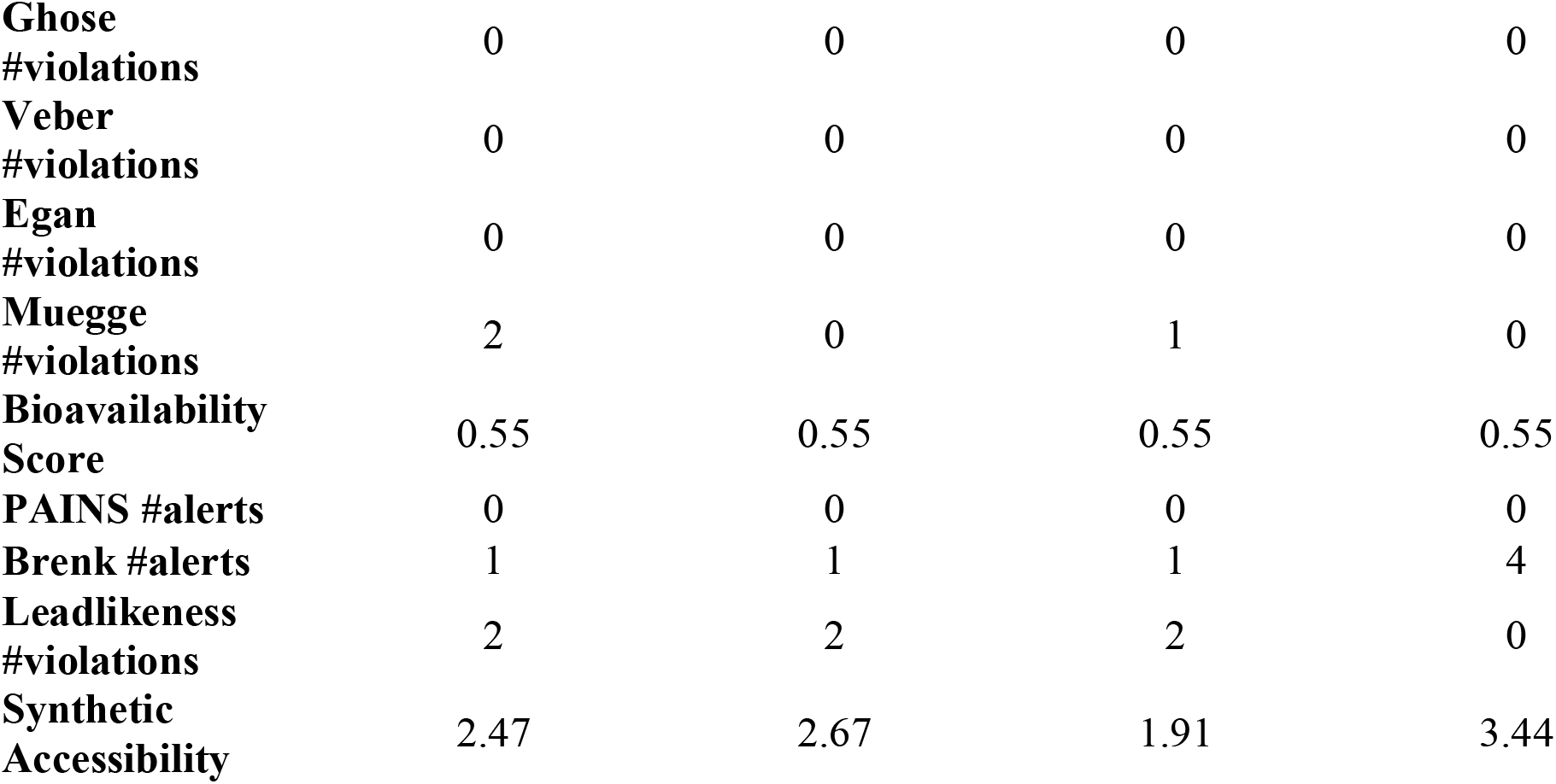
SwissADME Output for TSA and Curcuphenol and New Analogues.

### Determining the Maximum Tolerable Dosages for Curcuphenol Synthetic Analogues

Flow cytometry was employed to show that P02 and P03 curcuphenol analogues could trigger MHC-I expression on the cell surface 48 hours post-treatment while maintaining minimal cytotoxicity (**Figure 3B**). To identify the highest tolerable dose for curcuphenol analogues P02-113 (11) and P03-97-1 (7), various concentrations were assessed for toxicity. Dosages began at 1.0 mg/kg and increased to 3.5 mg/kg for both substances, reaching a final concentration of 5.2 mg/kg. The solubility of the compounds was the primary constraint in this experiment, as 1% DMSO is the maximum allowed concentration for intraperitoneal injections. Given these restrictions, 5.2 mg/kg was the largest intraperitoneal dosage employed. For each concentration of both compounds, three mice were examined, totaling nine mice per compound. Mice were observed for 14 days, and no indications of cytotoxicity were detected. Following the 14-day period, the mice underwent necropsy. No observable signs of toxicity or irregularities were found for either compound at any dosage. As a result, 5.2 mg/kg was selected as the dosage for subsequent studies.

### Pharmacokinetics of P02-113 and P03-97-1-1 in tumour-bearing mouse model

Pharmacokinetics of PC-02-113 (11) and PC-03-97-1 (7) were monitored after intraperitoneal injection at different time points to determine the dosing regimen for treatment of mice. Time points were chosen based on the literature for TSA, a structurally similar compound. TSA is broken down within 5 to 60 minutes, possesses a half-life of slightly less than 10 minutes, and is undetectable after 24 hours (43). Although the analogues are structurally similar to each other, their metabolism was significantly different. PC-03-97-1 (7) was found in mouse plasma at a concentration of 30 ng/ml at 5 minutes, reaching approximately half this concentration at approximately 20 minutes based on the 10- and 30-minute time points. Conversely, PC-02-113 (11) was detected at a concentration of 0.4 ng/ml within 5 minutes and reduced to half that amount after 10 minutes. As a result, the half-life of PC-03-97-1 (7) was determined to be 15 minutes, while the half-life of PC-02-113 (11) was estimated to be under 5 minutes. Time points prior to 5 minutes were not possible due to time-limited ability to inject and draw blood from mice. Another limitation was that one mouse could not be used for multiple time points, as each time point required one mouse to obtain sufficient plasma for PK sampling. Both compounds were consistent in reaching undetectable concentrations in mouse plasma at 6 hours. Due to the high eliminations in the mouse plasma, which is effective upon daily dosing, as well as in limitation dosing regimens, daily treatment was chosen as treatment for the *in vivo* studies. (**Figure 4**).

**Figure 4:**
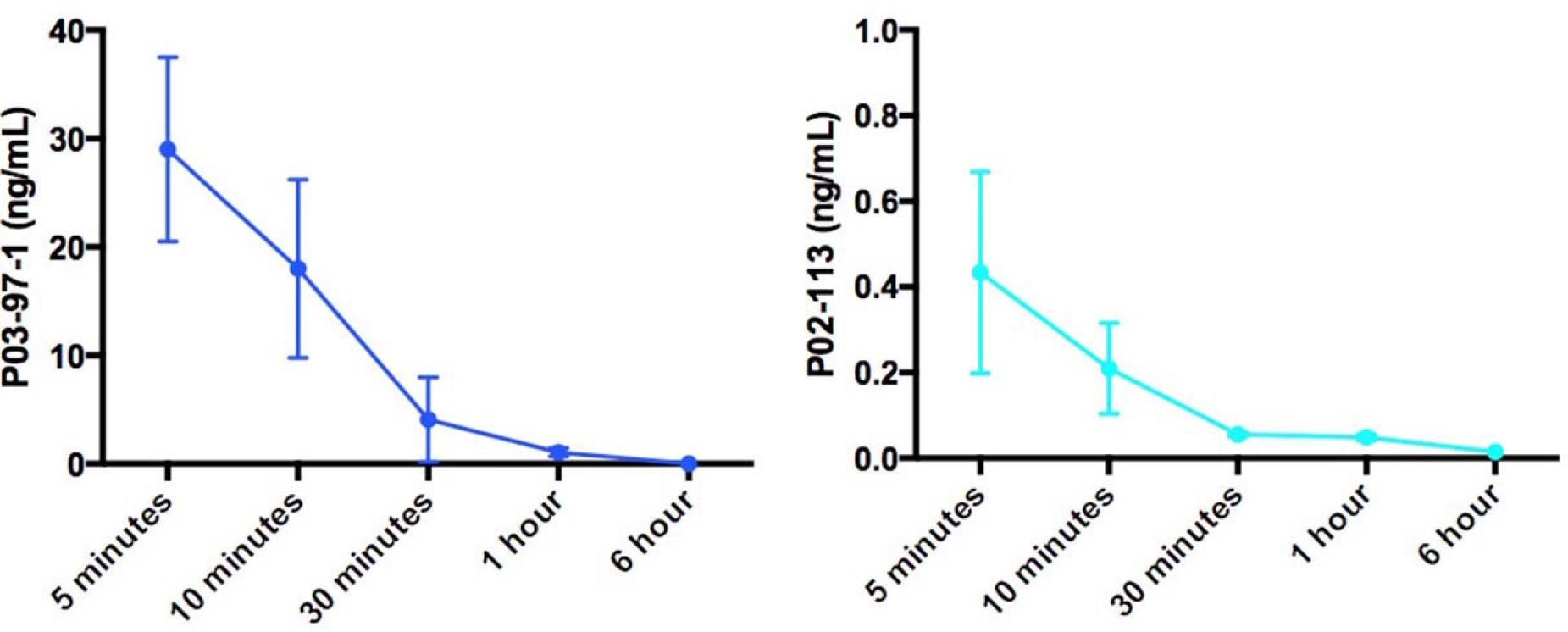
Pharmacokinetic analysis of P02-113 and P03-97-1: The average concentration of P02-113 and P03-97-1 in mouse plasma from three mice at each time point is shown with SEM. Female C57BL/6 mice, aged 6-8 weeks, received an i.p. injection of 5.2 mg/kg of P02-113 or P03-97-1, and blood was collected through cardiac puncture at different time points (n=3) after injection. Plasma was separated from blood and sent to The Metabolomics Innovation Center (TMIC) at the UVIC Genome BC Proteomics Centre for pharmacokinetic analysis, packed in dry ice.

### Evaluation of curcuphenol analogues to reduce the growth of metastatic tumours

Mice were injected subcutaneously (s.c.) in the right flank with 4×10^5^ A9 metastatic mouse lung cancer cells, and tumors were allowed to develop for 7 days (Figure 5A). Following the 7-day period, mice received daily treatments for 12 days with either TSA (positive control, 500 μg/kg), 1% DMSO (vehicle negative control), or one of the two experimental compounds, PC-02-113 (11) or PC-03-97-1 (7) (at 5.2 mg/kg). Body weight was recorded every 2-4 days, with no notable differences observed among the four groups (**Figure 5A**). Tumor sizes were assessed in all groups three times per week, and any mice that did not develop tumors during the study were excluded from tumor volume analysis. After administering treatments for 12 days, a statistically significant reduction in tumor volume was observed between the treated groups (PC-02-113 and PC-03-97-1) and the untreated group (1% DMSO), as established by a one-tailed t-test (p <0.0001) (**Figure 5B**).

**Figure 5.**
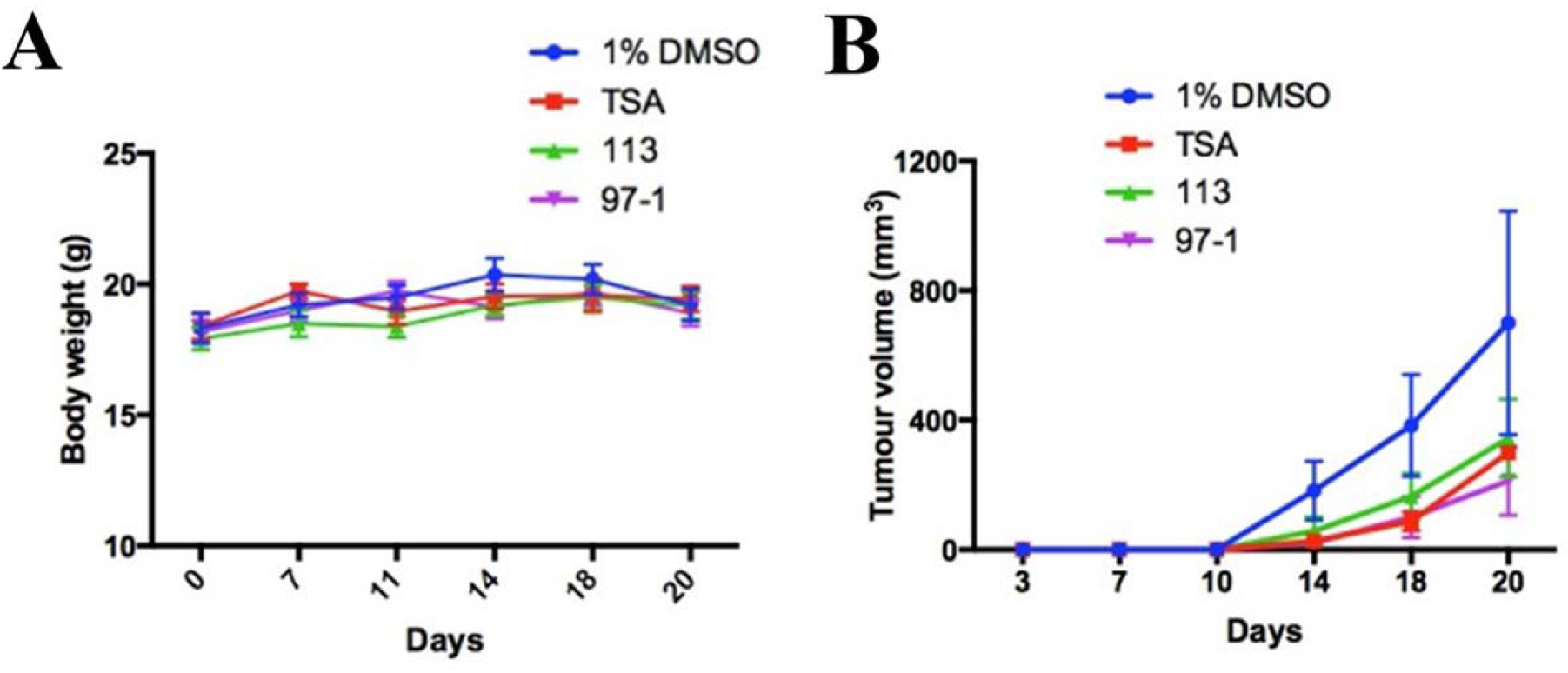
*In vivo* treatment with PC-02-113 or P03-97-1 hinders the growth of APM-deficient metastatic A9 lung cancer cells. A9 cells were injected subcutaneously into the right flank. After seven days, mice were divided into four treatment groups: (a) 1% DMSO vehicle negative control, (b) TSA (0.5mg/kg), (c) P02-113 (5.2mg/kg), and (d) P03-97-1 (5.2mg/kg). After 12 days of treatment, tumors were excised and analyzed for anti-CD4+ (APC) and anti-CD8+ (PE-Cy7) infiltration using flow cytometry. A) Body weights remained unaffected by the compounds, suggesting no toxicity. B) TSA, P02-113, and P03-97-1 significantly reduced tumor growth in comparison to the DMSO control established by a one-tailed t-test (p <0.0001).

### Enhancement of Tumour infiltrating lymphocytes by curcuphenol analogues

The two compounds, PC-02-113 (11) and PC-03-97-1 (7), exhibited remarkable anti-cancer therapeutic potential both in vitro and in vivo. For in vitro experiments, 1% DMSO served as a negative control. The ability of these compounds to induce both TAP-1 and MHC-I expression makes them excellent candidates for human clinical trials, as a decrease in the expression of these proteins is a characteristic of some cancer types. To gain a deeper understanding of the anti-tumor mechanisms of these compounds, tumors were also examined for endpoint T-cell infiltration. Tumors were assessed using CD4+ (APC) and CD8+ (PE-Cy7) T cell line cytometry (**Figure 6**). Intriguingly, CD8+ T-cell infiltration mirrored the pattern observed in tumor burden. TSA and PC-03-97-1 (7) exhibited the highest CD8+ penetration, followed by PC-02-113 (11) and vehicles alone. In terms of CD4+ penetration, no significant differences were observed among the groups. These findings suggest that PC-03-97-1 (7) acts as a more potent in vivo immunostimulant and demonstrates the most significant reduction in tumor burden, indicating that future research should concentrate on optimizing the structure of P03-97-1 (**Figure 6**).

**Figure 6.**
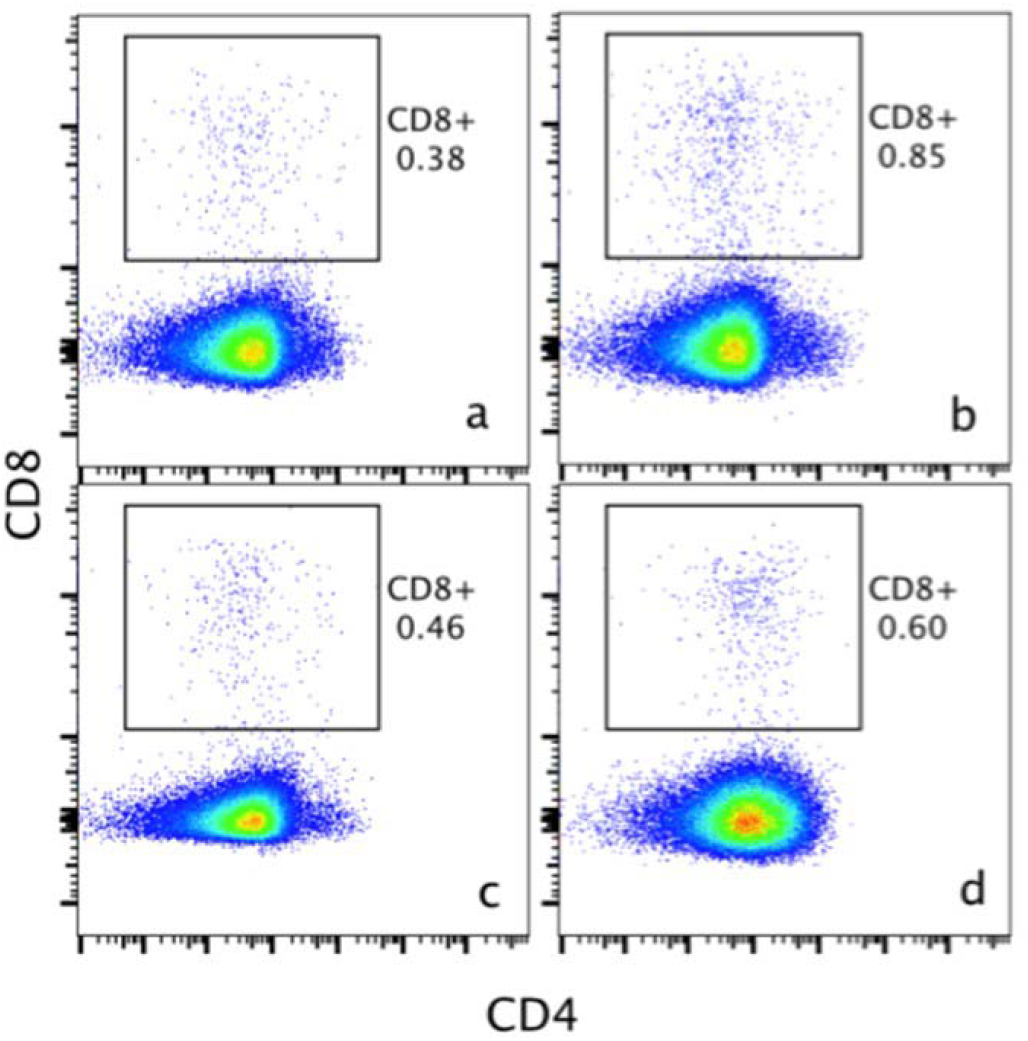
Evaluation of T cell infiltration in metastatic tumors in vivo. C57BL/6 mice were injected subcutaneously in the right flank with 5 x 104 APM-deficient metastatic A9 lung cancer cells. Seven days post-injection, mice were assigned to four treatment groups: vehicle (a), TSA (0.5mg/kg) (b), P02-113 (5.2mg/kg) (c), or P03-97-1 (5.2mg/kg) (d). After 12 days of treatment, tumors were excised and assessed for anti-CD4+ (APC) and anti-CD8+ (PE-Cy7) infiltration using flow cytometry.

## Discussion

Relatively few therapeutic modalities that reverse immune-escape or immune-editing have been developed. In this context, as a starting point, natural products provide an excellent source of novel anti-cancer compounds, with extracts that can be obtained from conventional food ingredients such as spices and herbs. Although spices are often considered as culinary staples, many spices have been identified with medicinal properties, including cumin, turmeric, saffron, green and black tea, and flax seeds containing curcumin. (44, 45, 46, 47, 48). Thus, examining extracts from natural sources remain a potentially important resource of new therapeutics, including those that may reverse immune-escape, and can reduce cancer growth and metastasis.

To contextualize the importance of targeting the reversal of immune-escape in cancer, Stutman (1974) for a time, had essentially extinguished the concept of T-lymphocytes mediating adaptive immune surveillance by purportedly showing that there was no difference between the growth of tumours in athymic nude mice missing T-cells versus wild-type animals (49, 50). Unfortunately, it appears Stutman was unaware while executing these studies that they were conducted with tumours that lacked APM capabilities and were therefore invisible to the host adaptive immune system and the presence of absence of T lymphocytes would make no difference in the context of immune escape where the tumours are selected by the immune system to become “invisible” to the T lymphocytes that negatively selected them. Since then, these conclusions have been formally refuted by Alimonti *et al*. (2000) (21) directly revisiting Stutman’s studies and conducting a parallel study to examine the growth of APM-competent and incompetent (immune-escape), chemically-induced tumours in athymic nude mice lacking only T-lymphocytes versus wild-type animals (21). This study demonstrated conclusively, and likely for the first time, that T-lymphocytes are required for immune surveillance, as is the coincident expression of functional APM and MHC-I in a tumour; setting forth the rules of immune engagement with tumours and setting the stage for engaging the power of T cells in cancer immunotherapies. Subsequently, Shankran et al (2002) confirmed the observations previously reported by Alimonti et al (2000) (21) in immune incompetent Rag2 knock-out (Rag2^-/-^) mice, which have a combined immune defect halting the development of T-cells, B cells and NK T-cells, and also in Stat-1-/- mice, which lack the IFN-γ receptor gene, and demonstrated both these mice lines also develop more chemically-induced sarcomas quicker than wild-type mice (22). Shankran et al (2001) re-coined the process of “immune-escape” (21) and referred to it as “immune-editing” (22). Thus, the selective pressure of immune surveillance on genetically unstable tumour populations yields tumours that have lost expression of APM components which results in the reduced assembly of functional major histocompatibility complex (MHC or HLA) molecules (13, 21, 38, 39). This phenotype is now commonly known to be associated with immune-escape that is linked to metastatic cancers (21, 37). For example, several types of cancers (29, 30, 31, 32, 33, 34, 35, 36) exhibiting APM deficits exhibit a clear link between human leukocyte antigen (HLA) down-regulation and poor prognosis (26, 27, 28). Dependent upon the specific tumour type, the loss of the APM components and functional MHC-I molecules with an immune-escape may appear in up to 90% of patients and has been linked to tumour aggressiveness and increased metastatic potential (26, 27, 28). Thus, in practice most immunotherapies that target metastatic cancers have to overcome immune-escape mechanisms coincidently with boost ant-cancer cell-mediated immune responses.

Here, curcuphenol, a component of turmeric used in curry spices, was identified as an active compound that induces the expression of TAP-1 and MHC-I molecules (APM components) in metastatic tumours. In this regard, curcuphenol, occurs naturally as one of two enantiomers: S- (+) and R- (-) curcuphenol (51, 52, 53, 54, 55, 56, 57, 58), however in the applications of interest only the S- (+) of curcuphenol is biologically active. Therefore, a pharmacopoeia of analogues of curcuphenol were synthesized in this study with the aim of finding more effective analogues. Curcuminoids are generally insoluble in water and we used simple chemical synthesis methods to create water soluble, an achiral curcuphenol analogue PC-02-113 (11) and a racemic analogue PC-03097-1 (7) that are soluble in both PBS and water and have enhanced chemical characteristics and biological performance, and we subsequently demonstrated their ability to enhance the expression of APM components which results in the reversal of the immune-escape phenotype in metastatic tumours. Clearly, these compounds have advantages in solubility and ease of synthesis and can be used as singular molecular entities rather than plant or animal extracts that contain many, sometimes counteractive chemical entities. We subsequently demonstrated their ability to reverse the immune-escape phenotype in metastatic tumours by enhancing the expression of APM components. The two enhanced curcuphenol-based analogues, PC-02-113 (11) and PC03-97-1 (7), that were further examined are well-tolerated *in vivo*, with no observable toxicity in animals at the examined dosages. When administered to metastatic tumor-bearing mice, these compounds led to a significant decrease in average tumour size. Overall, PC-03-97-1 (7) demonstrates a more potent effect in vivo, making it a promising candidate for future combination therapies. It has the potential to induce MHC-I molecule expression and improve survival rates for cancer patients with immune-escape phenotypes due to reduced APM levels. However, optimization of the dosage of PC-03-97-1 (7) and further chemical modification of the scaffold to increase its plasma half-life are necessary as it has been shown to have a high clearance rate from the plasma of rats as it became undetectable after six hours. This may be overcome by delivery in formulations such as lipid nanoparticles.

Studies conducted *in vitro* indicate that curcuphenol reverses the MHC-I immune-escape of metastatic tumours and thereby revealing the tumour to immune recognition and enhancing effector CD8+ T lymphocytes detection of metastatic tumours. In our studies, no increase in the frequency of CD4+ tumour infiltrating T lymphocytes was observed though it is conceivable that the activation and functional status of these cells may have changed in curcuphenol-analogue treated animals. However, a 2-fold increase in frequency of CD8+ tumour infiltrating T lymphocytes was observed. Despite this, it is difficult to determine from the literature the functional significance of a 2-fold increase in CD8+ tumour infiltrating T lymphocytes. It could suggest either the metastatic tumour growth inhibiting effect we observe could be due to this increase in CD8+ tumour infiltrating T lymphocytes or perhaps a systemic increases in tumour specific T lymphocytes, alternatively the effect we observe is due to the added contribution of other leukocyte population we are not measuring, such as natural killer cells and dendritic cells, or that some of the growth inhibiting effect is due to another mechanism that requires discovery. Future lines of investigation will be needed to resolve this question.

Overall, the fact that tumours may have acquired the ability to become ‘invisible’ to CTLs and may also become refractory to emerging cell-based immuno-therapeutics on utilizing T lymphocytes and immune checkpoint blockage inhibitors that seek to enhance T lymphocyte cytolysis against tumours is profoundly ignored in most cancer immunotherapies. Currently, only 15–30% of patients respond to existing immunotherapies (14, 59), and hence, discovering new therapeutic candidates that overcome immune-escape and augment the evolving immunotherapy modalities should be a priority. Therefore, combination therapies where the addition of drugs targeting the increased expression of the APM may be key in the future (10, 12, 38).

In conclusion, natural sources of curcuminoid molecules found in traditional spices such as turmeric and curcumin, have been used medicinally for perhaps thousands of years. However, this is the first studies that pinpoints reversing immune-escape mechanisms to explain the anti-cancer action of unique bio-active molecular entities derived from these extracts. In the greater sense, this study serves to highlight the medicinal value of common components of foods and establishes a new function for curcuminoid-containing preparations in promoting immunity to cancers.

## Experimental Procedures

### Experimental procedures for the synthesis of racemic curcuphenol and analogues

The general synthetic scheme used to synthesize the curcuphenol analogs PC-03-97-1 (**7**), PC-03-97-2 (**8**), PC-03-93 (**9**), PC-03-99 (**10**), PC-02-113 (**11**) and racemic curcuphenol (**12**) is shown in Figure 3 Panel A. The specific reaction details for the synthesis of PC-03-97-1 (**7**) and PC-03-97-2 (**8**) starting from aldehyde **1** are given below. The exact same reaction conditions were used to make racemic curcuphenol (**12**) and the analogues **9, 10** and **11** from starting materials **13**, **14**, **15**, and **19**, respectively, so the specific details of those syntheses have not been repeated here.

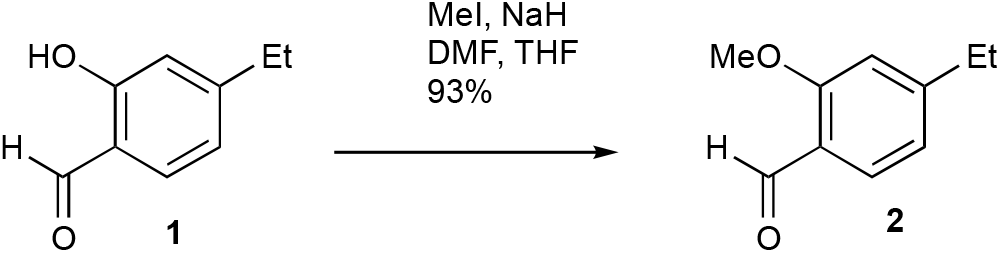

A suspension of NaH (172.6 mg, 60% in mineral oil, 4.32 mmol) in a DMF/THF mixture (5.4 mL, 4:1 v/v) was slowly combined with a solution containing 2-hydroxy-4-methoxybenzaldehyde (1) (546.6 mg, 3.60 mmol) and MeI (0.46 mL, 7.32 mmol) in THF (3.6 mL) at 0 °C. The cooling bath remained in place but was not replenished, and the mixture stirred for 18 hours. The mixture was then diluted with Et2O and washed with H2O. The organic extract was dried over MgSO4 and the solvent removed under vacuum. The resulting residue was purified through flash chromatography (silica gel, step gradient from 5:100 EtOAc/hexanes to 15:100 EtOAc/hexanes), yielding compound 2 (552.5 mg, 93%) as a white solid. The 1H NMR (400 MHz, CDCl3) showed δ 10.28 (s, 1H), 7.79 (d, J = 8.8 Hz, 1H), 6.53 (dd, J = 1.2, 8.4 Hz, 1H), 6.43 (d, J = 2.0 Hz, 1H), 3.89 (s, 3H), and 3.86 (s, 3H). The 13C NMR (100 MHz, CDCl3) presented δ 188.5, 166.4, 163.8, 130.9, 119.2, 106.0, 98.1, 55.81, and 55.79.

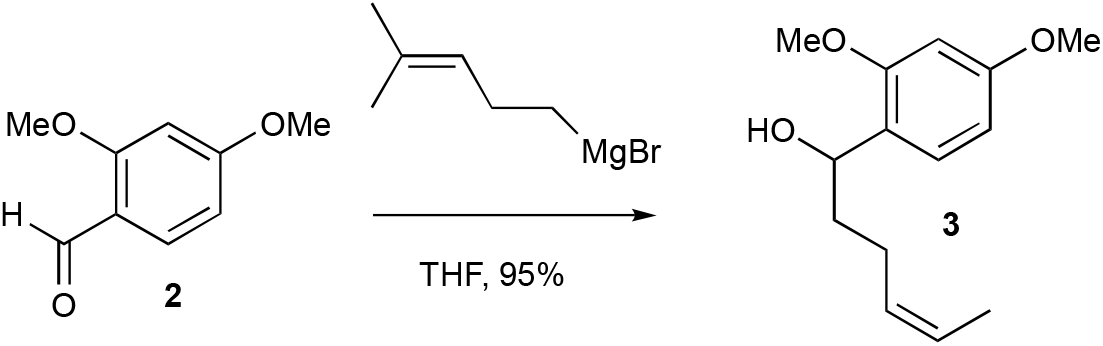

A mixture of Mg turnings (189.4 mg, 7.89 mmol) and a small amount of I2 in Et2O (1.0 mL) was combined with several drops of a 5-bromo-2-methyl-2-pentene (0.88 mL, 6.57 mmol) solution in Et2O (4.8 mL). After stirring for a few minutes, the yellow solution turned colorless, and the bromide solution was gradually added over an hour. The reaction mixture was then stirred under reflux for 1 hour. A freshly prepared Grignard reagent was added to a solution of compound 2 (272.3 mg, 1.64 mmol) in THF (8 mL) at 0 °C. The cooling bath was left in place but not recharged, and the mixture was stirred for 18 hours. The reaction was quenched with saturated aqueous NH4Cl solution and extracted with EtOAc three times. The combined organic extracts were washed with brine, dried over MgSO4, and evaporated under vacuum. The residue was purified by flash chromatography (silica gel, step gradient from 5:100 EtOAc/hexanes to 15:100 EtOAc/hexanes) to yield compound 3 (389.6 mg, 95%) as a colorless oil. The 1H NMR (400 MHz, CDCl3) showed δ 7.19 (d, J = 8.4 Hz, 1H), 6.44-6.48 (m, 2H), 5.15 (tt, J = 1.2, 7.2 Hz, 1H), 4.80 (t, J = 6.4 Hz, 1H), 3.81 (s, 3H), 3.79 (s, 3H), 2.53 (bs, 1H), 1.98-2.16 (m, 2H), 1.72-1.88 (m, 2H), 1.69 (s, 3H), and 1.59 (s, 3H). The 13C NMR (100 MHz, CDCl3) displayed δ 160.2, 157.9, 132.0, 127.8, 125.3, 124.4, 104.2, 98.8, 70.5, 55.5, 55.4, 37.4, 25.9, 25.0, and 17.9. HRESIMS [M + Na]+ m/z 273.1462 (calculated for C15H22O3Na, 273.1467).

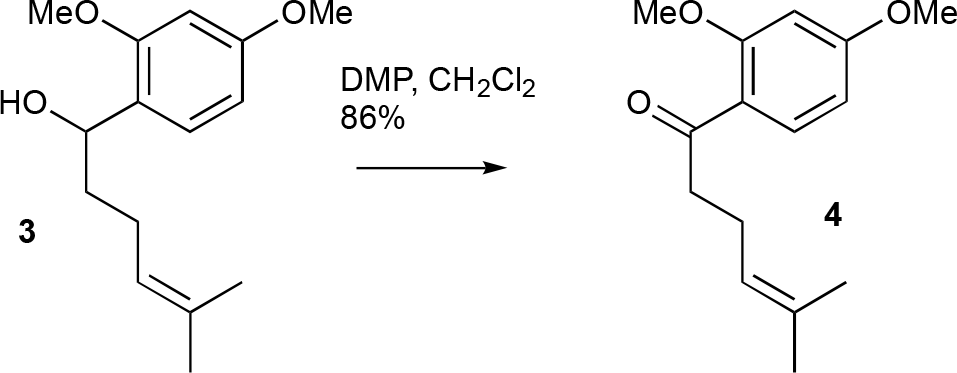

A solution of compound 3 (327.1 mg, 1.31 mmol) in CH2Cl2 (10 mL) was combined with DMP (714.9 mg, 1.64 mmol) at room temperature. After stirring for 30 minutes, TLC analysis confirmed the complete consumption of the starting material. Saturated aqueous NaHCO3 solution was then added, and the mixture was extracted with CH2Cl2 three times. The combined organic extracts were washed with brine, dried over MgSO4, and evaporated under vacuum. The residue was purified using flash chromatography (silica gel, step gradient from 0:100 EtOAc/hexanes to 8:100 EtOAc/hexanes), yielding compound 4 (280.3 mg, 86%) as a colorless oil. The 1H NMR (400 MHz, CDCl3) showed δ 7.79 (d, J = 8.8 Hz, 1H), 6.52 (dd, J = 2.0, 8.8 Hz, 1H), 6.45 (d, J = 2.0 Hz, 1H), 5.15 (dt, J = 1.6, 7.2 Hz, 1H), 3.88 (s, 3H), 3.85 (s, 3H), 2.95 (t, J = 7.6 Hz, 2H), 2.34 (q, J = 7.6 Hz, 2H), 1.68 (s, 3H), and 1.61 (s, 3H). The 13C NMR (100 MHz, CDCl3) displayed δ 200.5, 164.4, 160.9, 132.9, 132.2, 123.9, 121.5, 105.2, 98.5, 55.70, 55.65, 43.9, 25.9, 23.5, and 17.8. HRESIMS [M + Na]+ m/z 271.1309 (calculated for C15H20O3Na, 271.1310)..

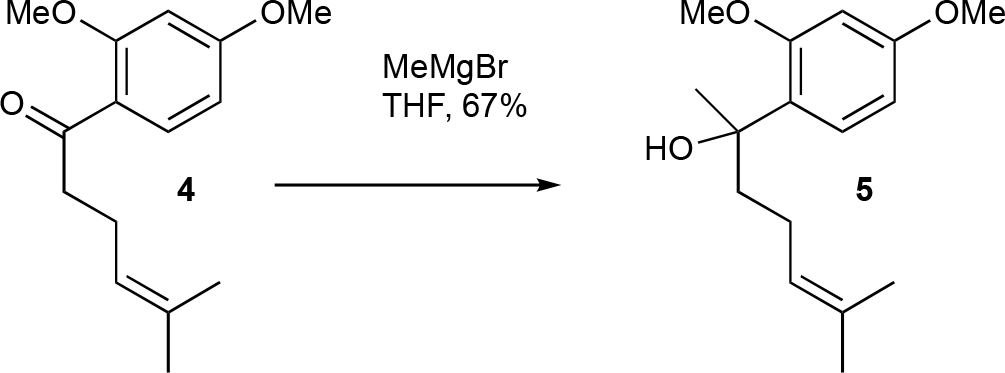

A solution of compound 4 (179.3 mg, 0.72 mmol) in THF (3 mL) had MeMgBr solution (0.32 mL, 3.0 M in Et2O, 0.96 mmol) slowly added at 0 °C. The mixture was stirred at room temperature for 2 hours. The reaction mixture was then cooled to 0 °C and quenched with saturated aqueous NH4Cl solution. The mixture was extracted with CH2Cl2 three times, and the combined organic extracts were washed with brine, dried over MgSO4, and evaporated under vacuum. The residue was purified using flash chromatography (silica gel, step gradient from 0:100 EtOAc/hexanes to 7:100 EtOAc/hexanes), yielding compound 5 (128.1 mg, 67%) as a colorless oil. The 1H NMR (400 MHz, CDCl3) showed δ 7.21 (d, J = 8.4 Hz, 1H), 6.49 (d, J = 2.4 Hz, 1H), 6.46 (dd, J = 2.4, 8.8 Hz, 1H), 5.08 (t, J = 6.8 Hz, 1H), 3.85 (s, 3H), 3.82 (bs, 1H), 3.80 (s, 3H), 1.79-2.01 (m, 4H), 1.65 (s, 3H), 1.54 (s, 3H), and 1.51 (s, 3H). The 13C NMR (100 MHz, CDCl3) displayed δ 159.8, 157.9, 131.6, 127.6, 127.5, 124.8, 104.1, 99.5, 75.0, 55.5, 42.3, 27.7, 25.8, 23.6, and 17.7. HRESIMS [M + Na]+ m/z 287.1619 (calculated for C16H24O3Na, 287.1623).

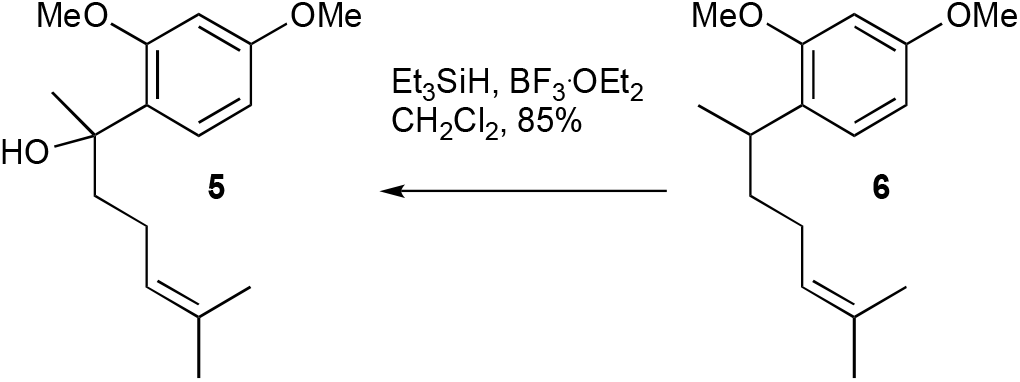

A solution of compound 5 (169.3 mg, 0.64 mmol) in CH2Cl2 (2 mL) had Et3SiH (0.13 mL, 0.81 mmol) added dropwise at -78 °C. After stirring for 10 minutes, BF3.OEt2 (0.12 mL, 0.97 mmol) was added dropwise, and the mixture was stirred for an additional hour at -78 °C. The mixture was then diluted with CH2Cl2 and washed with saturated aqueous NaHCO3 solution and water until neutral. The organic extract was dried over MgSO4 and evaporated under vacuum. The residue was purified using flash chromatography (silica gel, step gradient from 0:100 EtOAc/hexanes to 1:100 EtOAc/hexanes), yielding compound 6 (135.1 mg, 85%) as a colorless oil. The 1H NMR (400 MHz, CDCl3) showed δ 7.06 (d, J = 8.0 Hz, 1H), 6.45-6.48 (m, 2H), 5.10-5.15 (m, 1H), 3.802 (s, 3H), 3.797 (s, 3H), 3.10 (sixt, J = 7.2 Hz, 1H), 1.83-1.97 (m, 2H), and 1.47-1.68 (m, 8H). The 13C NMR (100 MHz, CDCl3) displayed δ 158.8, 158.2, 131.3, 128.6, 127.3, 125.1, 104.2, 98.7, 55.54, 55.48, 37.5, 31.5, 26.5, 25.9, 21.4, and 17.8. HRESIMS [M + H]+ m/z 249.1854 (calculated for C16H25O2, 249.1855).

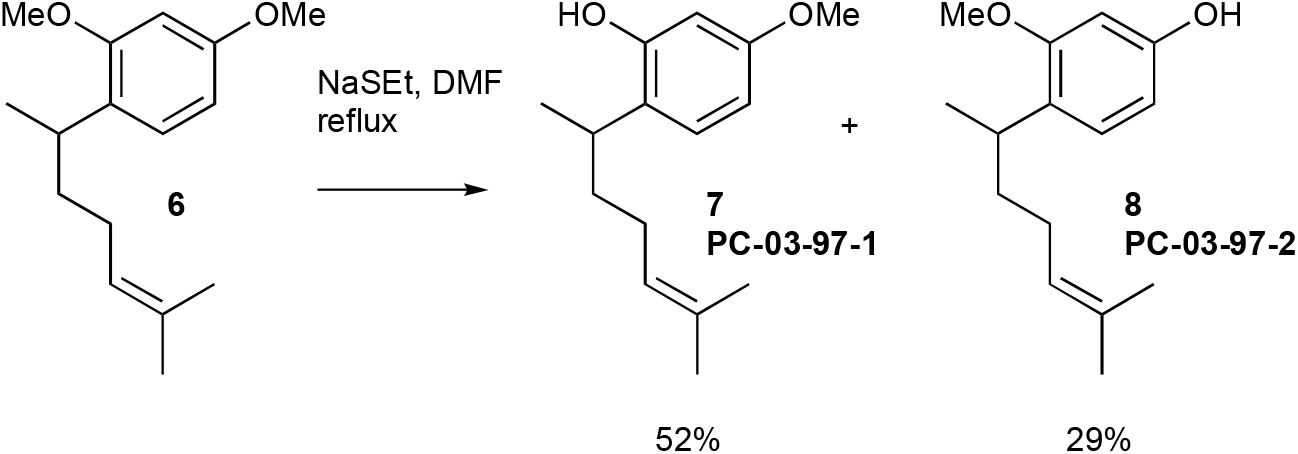

NaSEt (489.9 mg, 5.24 mmol) was mixed with DMF (2 mL) at 0 °C, then warmed to room temperature. A solution of compound 6 (110.0 mg, 0.44 mmol) in DMF (1 mL) was added, and the mixture was stirred under reflux for 3 hours. It was then cooled to 0 °C, and 10% HCl (∼3 mL) and CH2Cl2 (∼15 mL) were added. The organic layer was washed twice with water, dried over MgSO4, and evaporated under vacuum. The residue was purified using flash chromatography (silica gel, step gradient from 0:100 EtOAc/hexanes to 10:100 EtOAc/hexanes), yielding compound 7 (PC-03-97-1) (54.2 mg, 52%) as a colorless oil and compound 8 (PC-03-97-2) (30.4 mg, 29%) as a light yellow oil.

For isomer 7: The 1H NMR (600 MHz, CDCl3) showed δ 7.05 (d, J = 8.4 Hz, 1H), 6.49 (dd, J = 2.4, 8.4 Hz, 1H), 6.37 (d, J = 2.4 Hz, 1H), 5.14 (t, J = 7.2 Hz, 1H), 4.93 (bs, 1H), 3.77 (s, 3H), 2.92 (sixt, J = 7.2 Hz, 1H), 1.90-1.98 (m, 2H), 1.70 (s, 3H), 1.55-1.69 (m, 2H), 1.55 (s, 3H), and 1.23 (d, J = 7.2 Hz, 3H). The 13C NMR (150 MHz, CDCl3) displayed δ 158.6, 154.1, 132.4, 127.7, 125.5, 124.8, 106.5, 101.9, 55.5, 37.6, 31.3, 26.2, 25.9, 21.4, and 17.9. HRESIMS [M - H]- m/z 233.1543 (calculated for C15H21O2, 233.1542).

For isomer 8: The 1H NMR (600 MHz, CDCl3) showed δ 6.99 (d, J = 8.4 Hz, 1H), 6.39 (s, 1H), 6.38 (d, J = 7.8 Hz, 1H), 5.11 (t, J = 6.6 Hz, 1H), 4.81 (bs, 1H), 3.77 (s, 3H), 3.07 (sixt, J = 7.2 Hz, 1H), 1.82-1.96 (m, 2H), 1.47-1.69 (m, 8H), and 1.15 (d, J = 6.6 Hz, 3H). The 13C NMR (150 MHz, CDCl3) displayed δ 158.

## *In vivo* efficacy studies

### A9 Cell Culture

The A9 metastatic cell line is derived from the TC-1 murine lung carcinoma cell line, which itself originated from primary lung epithelial cells of a C57BL/6 mouse. The cells were immortalized using the LXSN16 retrovirus vector carrying Human Papillomavirus E6/E7 oncogenes and later transformed with the pVEJB plasmid expressing the activated human H-Ras oncogene. The A9 cell line was generated in vivo after an immunization approach in animals with the original TC-1 parental cells, promoting the selection of clones with increased immune resistance. Unlike the parental TC-1 cells, which exhibit high expression of TAP 1 and MHC I, A9 cells have nearly undetectable MHC I levels. A9 cells were grown in Dulbecco’s Modified Eagle’s Medium (DMEM; Gibco) supplemented with 10% fetal bovine serum (FBS; Gibco), 100 U/mL penicillin-streptomycin (Gibco), and maintained at 37 °C in a humidified atmosphere containing 5% CO2.

### Induction of MHC-I and TAP-1 Expression in A9 cells via Small Molecule Treatment

A9 cells were plated at a density of 3×10^5^ onto a 6-well plate in two mL of DMEM. Twenty-four hours after plating, cells were subjected to one of the following treatments for forty-eight hours:1% Dimethyl Sulfoxide (Sigma, #276855) in DMEM (1% DMSO) as a vehicle control, 150 nmol of Curcuphenol (kindly donated by Dr. Raymond Anderson) or 40 ng of Recombinant Mouse Interferon-γ (IFNγ; R&D Systems, 485-MI-100) as a positive control. Curcuphenol stock was dissolved in DMSO. All compounds were diluted in 1% DMSO.

### Reverse Transcription and Real-Time PCR

RNA was isolated using the Purelink^TM^ RNA Mini Kit (Invitrogen, #12183-025), followed by DNase I (Ambion, #AM2222) treatment. RNA was then reverse transcribed into cDNA using the SuperScript^®^ III First-Strand Synthesis System for RT-PCR (Invitrogen, #18080-051). Real-Time PCR (RT-PCR) was done using a 7500 Fast Real-Time PCR system (Applied Biosystems) with the following parameters: 40 cycles (95 °C denaturing 15 seconds, 60 °C annealing for 1 minute).

#### Maximum Tolerated Dose (MTD)

The highest tolerable doses for the selected compounds, P02-113 and P03-97-1, were evaluated in vivo. C57Bl/6 mice were given intraperitoneal (i.p.) injections of the compounds at three different concentrations: 1.0 mg/kg (n=3), 3.5 mg/kg (n=3), and 5.2 mg/kg (n=3). Mice were monitored for 14 days for any clinical signs of toxicity, and a necropsy was performed at the end of the study. The highest MTD dose did not cause any adverse effects and was utilized for further experiments.

#### Pharmacokinetic Study

Compounds PC-02-113 and PC-03-97-1 were assessed at three time points after an i.p. injection at a concentration of 5.2 mg/kg. Three mice were used for each time point, totaling nine mice per compound. Time points were chosen based on the compounds’ similarity to TSA, an HDACi known to enhance TAP-1 and MHC-I expression in metastatic tumors and to have a high metabolism rate. The selected time points were 5 minutes, 10 minutes, 1 hour, and 6 hours.

#### Treatment of Tumor-Bearing Mice with Identified Compounds

4×105 metastatic A9 cells suspended in HBSS were subcutaneously (s.c.) transplanted into the right flank of 32 eight-week-old female C57Bl/6 mice. Starting seven days after tumor injection, mice in each tumor group were treated daily by i.p. injection with either one of the identified compounds (n=8 for each compound), TSA positive control (n=8), or vehicle alone (n=8) for two weeks. Body weight and tumors were measured every 2-4 days, with more frequent measurements as tumor size increased. Tumor volume was calculated as follows: tumor volume = length x width^2. Tumor growth rate assessment methods were based on previously described techniques (17).

#### Survival Curves

Survival for mice with A9 tumors was determined based on the overall weight of the mouse and tumor volume. Mice were euthanized if they lost 20% of their initial weight or tumors grew larger than 1 cm3, in accordance with animal ethics guidelines.

### Evaluation of MHC-I Surface Expression by Flow Cytometry

Primary TC-1 tumor or metastatic A9 tumor cells, as shown in Figure 1, were plated in 6-well plates at a concentration of 10,000 cells per well in a 2 mL volume. Cells were treated with the indicated compound concentrations the following day and incubated for 48 hours at 37°C. After incubation, cells were trypsinized, washed, and stained with APC-conjugated anti-mouse MHC-I (specifically anti-H-2Kb) antibody (cat. 141605; clone 25-D1.16; Biolegend) and analyzed by flow cytometry. Primary TC-1 tumor cells served as a positive control for surface MHC-I expression, while vehicle alone (1% DMSO) was used as a negative control. Supplementary Table 1 displays the antibody panel used for FACS during the mouse tumor trial, and Supplementary Table 2 lists the primer sequences used for RT-PCR.

### Statistical Analysis

Data were examined using R and Excel. A Student’s t-test was utilized to establish statistical significance between controlled test groups, with a p-value of ≤ 0.05 deemed significant.

## Acknowledgments

We would like to thank Dr Eliana Al Haddad for her editorial assistance. This work was supported by an Industrial Partnered Collaborative Research grant from the Canadian Institutes of Health Research (CIHR; MOP-102698) to WAJ; a grant from the Natural Sciences and Engineering Research Council (NSERC; RGPIN 869-13) to RJA; and TM was funded through a NSERC Bioinformatics CREATE Program co-hosted at the University of British Columbia and Simon Fraser University.

## Author contribution

Conception of the Project: WAJ

Designed research: LLN, SLSE, CGP, DEW, RJA, WAJ

Performed research: LLN, SLSE, SD, IS, LM, BAE, DEW, PC, KBC, EG

Analyzed data: LLN, SLSE, SD, IS, LM, CGP, BAE, TM, DEW, PC, NAL, RJA, KBC, EG, WAJ

Wrote paper: LLN, SLSE, RJA, WAJ

Edited paper: LLN, SLSE, CGP, TM, DEW, NAL, RJA, WAJ

## Data Availability

The data that support the findings of this study are available from the corresponding author, WAJ, upon reasonable request.

---

**Supplementary Table 1:** Antibody panel used for FACS during mouse tumour trial

| Target | Fluorophore (Anti-mouse) | Clone | Catalog # | Company | Ratio used |
| --- | --- | --- | --- | --- | --- |
| CD3E | APC-eFluor 780 | 145-2C11 | 47-0031-82 | Invitrogen | 1:200 |
| CD11C | eFluor 506 | N418 | 69-0114-82 | Invitrogen | 1:200 |
| CD4 | Alexafluor 700 | GK1.5 | 56-0041-82 | Invitrogen | 1:200 |
| CD69 | PE-Cyanine7 | H1.2 F3 | 25-0691-82 | Invitrogen | 1:200 |
| CD45R | FITC | RA3-6B2 | 11-0452-82 | Invitrogen | 1:200 |
| MHCII | eFluor 450 | AF6-120.1 | 48-5320-82 | Invitrogen | 1:200 |
| CD49B | APC | DX5 | 12-5971-82 | Invitrogen | 1:200 |
| CD11B | SuperBright 645 | M1/70 | 64-0112-82 | Invitrogen | 1:200 |
| CD62L | SuperBright 702 | MEL-14 | 67-0621-82 | Invitrogen | 1:200 |
| MHC Class I (H-2KB) | PE | AF6-88.5.5.3 | 12-5958-82 | Invitrogen | 1:200 |

**Supplementary Table 2.**
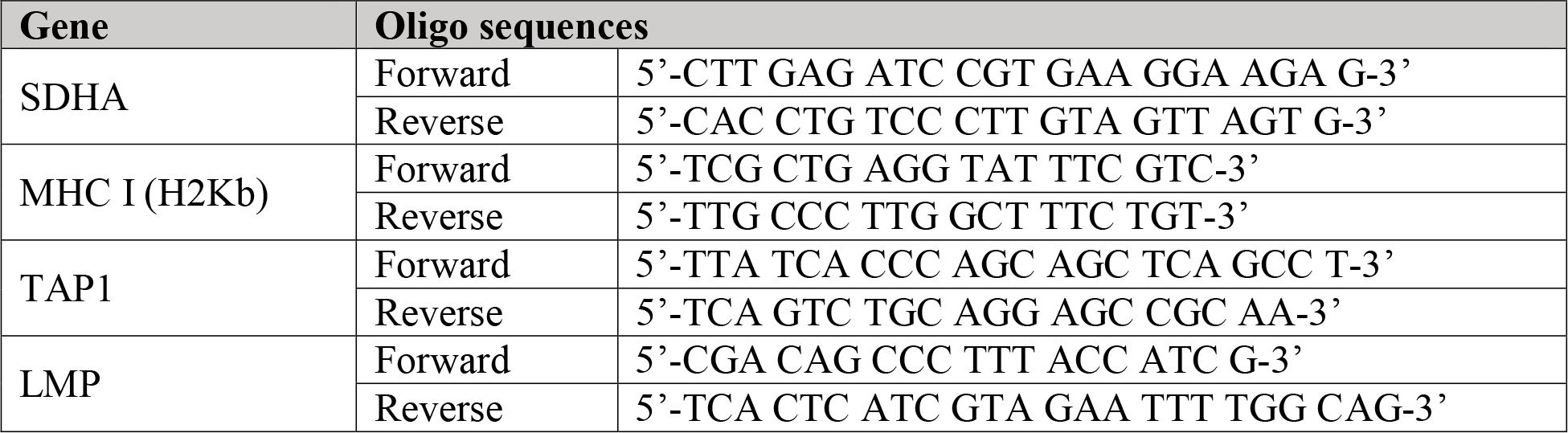
Primer sequences for RT-PCR

